# BondGraphs.jl: Composable energy-based modelling in systems biology

**DOI:** 10.1101/2023.04.23.537337

**Authors:** Joshua Forrest, Vijay Rajagopal, Michael PH Stumpf, Michael Pan

## Abstract

**Summary:** BondGraphs.jl is a Julia implementation of bond graphs. Bond graphs provide a modelling framework that describes energy flow through a physical system and by construction enforce thermodynamic constraints. The framework is widely used in engineering and has recently been shown to be a powerful approach for modelling biology. Models are mutable, hierarchical, multi-scale, multi-physics, and BondGraphs.jl is compatible with the Julia modelling ecosystem.

**Availability and Implementation:** BondGraphs.jl is freely available under the MIT license. Source code and documentation can be found at https://github.com/jedforrest/BondGraphs.jl.

**Supplementary information:** Supplementary data are available at *Bioinformatics* online.

## 1. Introduction

The last decade has seen an explosion of experimental methods that simultaneously measure many aspects of biological processes at multiple temporal and spatial scales. Extracting biological mechanisms from this big-data explosion requires modular, interoperable and computationally efficient software in systems biology (Rajagopal et al., 2022).

Bond graph modelling is a promising approach to create computational models for large-scale biological data analysis. This mathematical framework has been used to model many biological systems, as well as mechanical, electrical, and chemical engineering systems (Gawthrop and Bevan, 2007; Gawthrop and Pan, 2022).

Bond graph models are particularly advantageous for systems biologists and mathematical modellers. Bond graph based models of biological networks are biophysics-based and are scalable to large networks of interactions; they satisfy the laws of thermodynamics implicitly, and therefore they enable efficient construction of models that couple interactions between different sub-systems within a biological process. Recent biological bond graph models include mitochondrial respiration (Gawthrop et al., 2020), two component bacterial systems (Forrest et al., 2023), and vascular blood flow (Safaei et al., 2018).

Here we present **BondGraphs.jl**, a Julia implementation of the bond graph modelling framework. The package enables a wide range of analyses to be conducted on physical systems, including the derivation of differential equations and simulation. Graphs and equations are built with a high level symbolic interface that integrates closely with the Julia modelling ecosystem.

## 2. Features

The most pertinent features are listed below. We refer to the documentation^1^ for a full list of features and examples, and to the supplementary material for the basic principles of bond graph modelling.

### 2.1. Graphical construction and modification of models

One of the strengths of bond graphs is that they are semantic representations of models, much like an electrical circuit or free body diagram. Physical systems are represented as graphs, and can be manipulated in many of the same ways.

BondGraphs.jl constructs a `BondGraph` object using `Components` and `Junctions` (graph vertices) and `Bonds` (graph edges).

Each `Component` encodes acausal constitutive relations equation that represent a physical element, event, or law; for example Ohm’s law of electrical resistance. `Junctions` are vertices that describe how components relate to each other according to conservation laws; for example Kirchhoff’s Voltage Law. `Bonds` are the graph edges that connect the vertices and represent energy flow or transfer between parts of the system.

Bond graphs can either be constructed one component at a time and connected together with bonds (using `add_node!` and `connect!`), or generated automatically using specialised algorithms (see Section 2.3).

Extra functions are included for modifying and composing multiple bond graph objects. Vertices (components or junctions) can be swapped in for another (`swap!`). This is useful for rapidly creating bond graphs of systems that are structurally similar but with modified equations, such as a different constitutive relation for a spring or resistor. For example, in a biochemical context one could replace standard mass-action rate laws with Michaelis-Menten rate laws.

Vertices can also be inserted between two already connected components (`insert_node!`), which is useful for adding a conservation law (with a junction) or a transformation to the variables.

When combining two separate bond graphs, there are two main approaches. The first is to take the two disconnected graphs and merge nodes representing the same object (`merge_nodes!`). This way the bond graph retains a ‘flat’ structure and system elements are not duplicated. Alternatively, bond graph can be nested inside a `BondGraphNode` component, which functions like an ordinary component but contains another bond graph. This creates a hierarchical bond graph that is a powerful tool for model organisation (Pan et al., 2021).

These operations enable high level semantic construction and modification of the model. Since all models are built and stored as a graph object, they can be traversed and manipulated using existing graph algorithms (see Section 2.4).

BondGraphs.jl includes a library of common bond graph component types. The default library includes a biochemical library for modelling chemical species and reaction networks. Additional custom components can be defined by the user and stored in the local library. See supplementary material for further details.

### 2.2. Symbolic generation of equations

Systems of differential equations are automatically generated from the bond graph. Stored equations are systematically composed using a computer algebra system (CAS) to automatically substitute, simplify, and differentiate equations. This produces a system of ordinary differential equations (ODEs) or differential algebraic equations (DAEs). These equations can be solved numerically or analysed analytically using other packages in the Julia ecosystem (Section 2.4). A modeller is not limited to differential equations, as they can derive other useful representations such as Hamiltonians, reaction fluxes, or power equations.

The benefit of this approach is that the model building and equation solving steps are separated. Abstractions of models and repeated motifs can be stored and reused for different modelling contexts (Gawthrop and Bevan, 2007; Pan et al., 2021). The parameters can be decided at the simulation stage, and the CAS simplification routines reduce numerical instability. With Julia arbitrary functions can be used as control inputs, as long as they return a single real-valued output.

### 2.3. Generating models from chemical reaction networks

An important feature for systems biologists is the chemical reaction network ⟶ bond graph ⟶ differential equation pipeline. BondGraphs.jl provides an interface for Catalyst.jl (Loman et al., 2022) which converts a reaction network into a bond graph. This enables rapid construction and analysis of models without needing to manipulate the graph structure or type equations directly (though this is still possible if desired). See Section 3.2 for an example. Existing models that were originally represented as chemical reaction networks can therefore be easily converted into bond graphs.

### 2.4. Integration with the Julia modelling ecosystem

Previous bond graph packages have largely been implemented through proprietary software languages such as MATLAB or Maple. More recently, the open source package BondGraphTools implemented the framework in Python (Cudmore et al., 2021). However, since bond graphs are implemented as specialised Python objects, it is not possible to use or study the graph-theoretic properties of the model, or to easily integrate with other modelling or analysis libraries.

By contrast, BondGraphs.jl is built purely in Julia. Julia is computationally fast and scalable while retaining easy-to-read syntax. It is therefore, we would argue, better positioned to address the grand challenges of systems biology (Roesch et al., 2023). The data types and functionality of BondGraphs.jl are built upon widely used packages, which gives users access to all their features. This also enables close integration with the Julia modelling ecosystem. `BondGraphs` are `AbstractGraphs` as defined in Graphs.jl (Fairbanks et al., 2021). With this we can use a library of graph operations, traversal algorithms, graph colouring, database storage, and plotting recipes (as seen in Figure 1). ODEs are generated and stored using ModelingToolkit.jl (Ma et al., 2022) and are solved with the parallelisable numerical solver library DifferentialEquations.jl (Rackauckas and Nie, 2017). Bridging BondGraphs and ModelingToolkit connects our bond graphs to many other useful libraries, including parameter estimation and optimisation, model storage and collection, plotting libraries, and reaction network interfaces (Section 2.3). This also makes the package code easier to maintain, and updates to other packages will automatically be available for use in BondGraphs.jl.

**Fig. 1:**
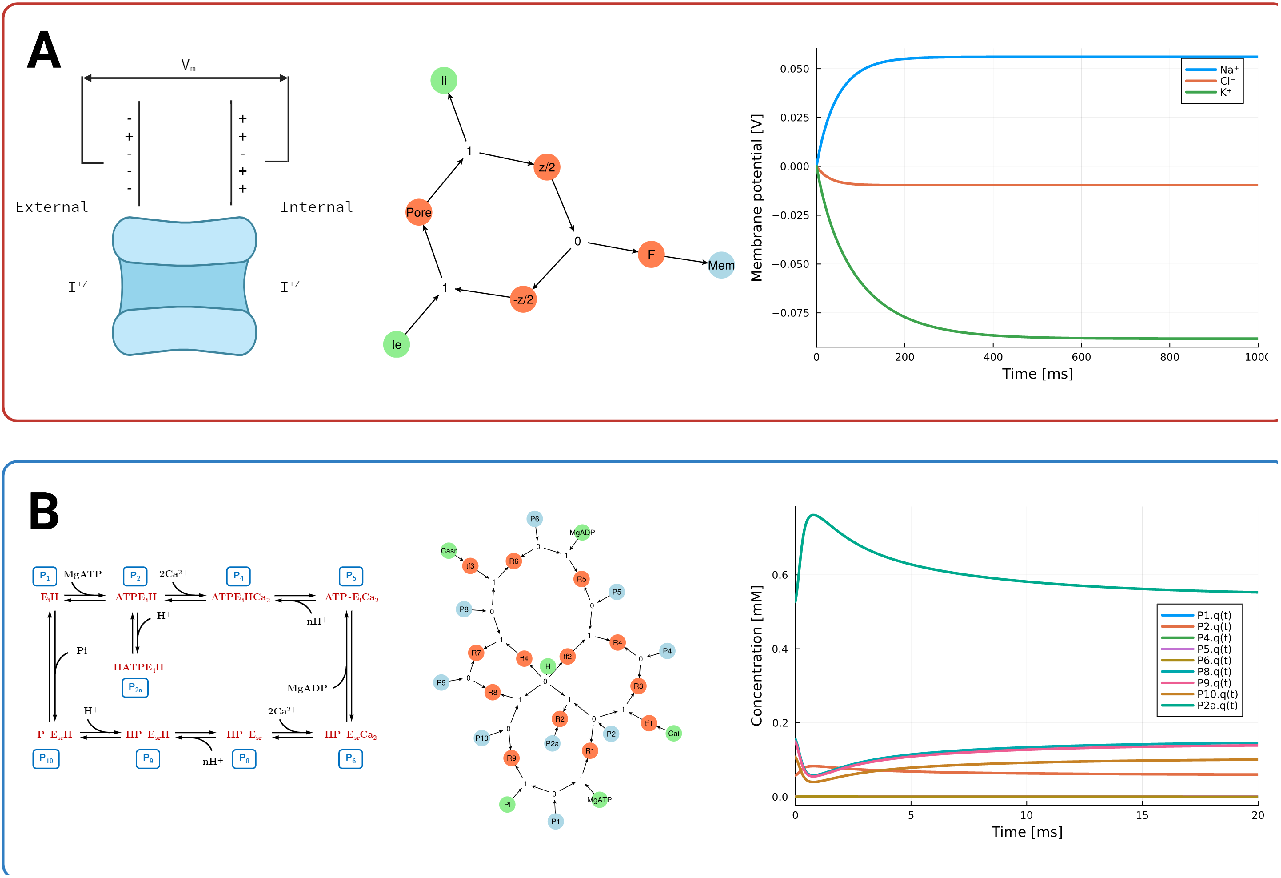
Two example physiological systems (left) alongside their bond graph formulation (centre) and steady state numerical solutions (right). **(A)** Ion pore channel (Cudmore et al., 2021) demonstrates modelling of a multiphysics (electro-chemical) system and repeated symbolic generation of models. **(B)** SERCA pump (Pan et al., 2020) demonstrates ease of scaling up biochemical bond graph modelling using the reaction network interface. Code used to generate the bond graphs and numerical solutions are available in the supplementary material. Diagrams created with BioRender.com.

## 3. Results

Here we demonstrate the features of BondGraphs.jl through two biological examples from literature. These two examples highlight several important features of both bond graphs and BondGraphs.jl. All code used to generate these plots and results can be found in the supplementary material.

### 3.1. Ion transport

Many biological systems are multi-physics in nature. Here we model an ion pore transport process, which couples electrical and chemical processes that drive movement of ions across cell membranes (Cudmore et al., 2021). The diagram, bond graph, and numerical solution are shown in Figure 1A.

Each vertex in the bond graph (middle column) represents a different physical element or law of the system. The membrane potential is an electrical capacitor (blue). The external and internal ion concentrations are constant sources of chemical energy (green). Orange vertices are two-port components that either ‘transform’ or ‘dissipate’ energy: the energy-dissipating ion transport reaction process (*pore*); the conversion of electrical to chemical potential (*F*); and the splitting of voltage and charge dependence across the two sides of the membrane (*z/2* and *-z/2*). The junctions (**0** and **1**) are the physical laws that enforce conservation of molar flow and chemical potential rate respectively. Bonds (edges) are the flow of energy between components, with the arrowhead indicating sign convention.

Components are constructed with parameter values and initial conditions, however these can be determined later. All nodes are added to the bond graph and connected with bonds in a graph-like manner. Once the bond graph is made, it can easily be reused with different values for different ions. Solutions for three ion transporters (Na^+^, K^+^, Cl^−^) are included in Figure 1A, right column. The steady state solutions follow the well-known Nernst equations seen in electrophysiology (Cudmore et al., 2021).

### 3.2 SERCA pump

Figure 1 shows a bond graph construction of the Sarco/Endoplasmic Reticulum Ca^2+^-ATPase (SERCA) pump, first presented in Tran et al. (2009) and subsequently represented as a bond graph in Pan et al. (2020). Reaction network and parameter values are taken from these papers. Figure 1B shows the reaction network schematic, bond graph, and numerical solution for species’ concentrations over time.

The colour coding follows the same convention as the previous example: blue vertices store chemical energy for one species; green vertices are sources of chemical energy (chemostats); orange vertices are reactions; junctions are physical conservation laws.

The SERCA bond graph model is generated using the reaction network interface (Section 2.3) with the following compact syntax:

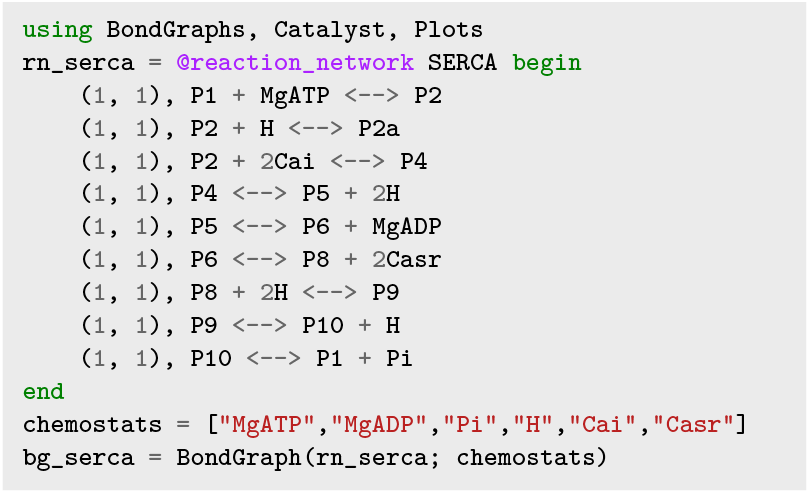

Note that the (1, 1) before each equation are placeholder reaction rates required for the catalyst interface.

This example highlights several features: (1) Model construction scales efficiently even for large complex reaction networks; (2) Bond graphs can be constructed from only the reactions network and interfaced at a high level without direct interaction with equations; (3) Parameters and initial conditions are determined separately and can include control variable equations; the input concentration of Ca^2+^ is set as a nonlinear “spike” function over time (see supplementary material for details).

## 4. Conclusion

BondGraphs.jl is a new powerful tool for large, multi-physics, biophysically constrained systems biology models. Bond graphs are a general framework beyond biology, however they have specific benefits to the systems biologist: (i) Models are constructed abstractly without worrying about equation implementation; (ii) Governing equations are derived automatically; (iii) The energy-based framework ensures that generated models are physically and thermodynamically compliant; (iv) BondGraphs.jl naturally integrates with other state-of-the-art tools in the Julia modelling ecosystem.

## Acknowledgements

JF was supported by the Melbourne Research Scholarship from the University of Melbourne. MP was supported by a Postdoctoral Research Fellowship from the School of Mathematics and Statistics at the University of Melbourne.

## Supplementary Material

### A. Bond graph theory

This section covers the basic principles of bond graph modelling. Table 1 summarises the variable and component analogies discussed in this section.

**Table 1.** Bond graph variables and analogous components for common physical domains.

|  | Bond Graph | Mechanical | Electrical | Chemical |
| --- | --- | --- | --- | --- |
| Quantity [$x$] | $x$ | [metres] | [coulombs] | [moles] |
| Effort [Joules/$x$] | $e$ | Force | Voltage | Chemical potential |
| Flow [$x$/second] | $f$ | Velocity | Current | Molar flow rate |
| Integrated Effort | $p = \int e dt$ | Momentum | Magnetic flux | $n/a$ |
| Integrated Flow | $q = \int f dt$ | Displacement | Charge | Concentration |
| Compliance | **C** | Spring | Capacitor | Species |
| Resistance | **R** | Damper | Resistor | Reaction |
| Impedance | **I** | Inertia | Inductor | $n/a$ |
| Transform | **TF** | Lever | Transformer | Stoichiometry |

Bond graphs represent all systems and reactions in terms of energy flow between elements of a system. Physical variables are described in terms of abstracted *efforts* (forces, voltages, chemical potentials) and *flows* (velocities, currents, molar flow rate). Efforts are always in terms of Joules per some quantity *x*, and flows are always *x* per second, so that *J/x · x/s* = *J/s*, or energy change over time. By abstracting these units, any of the above efforts and flows can be combined in a single model that conserves energy across all domains.

Bond graphs are built with collections of components. Components are generalised forms of physical laws, and typically represent a physical element of the system, such as a resistor, point mass, or chemical species. Each component stores a constitutive relation that describes the relationship between the energetic *effort* [Joules/*x*] and energy *flow* [*x*/second] in the component, where *x* is a measured quantity (displacement, charge, moles). BondGraphs.jl includes a library of common bond graph component types. These include the **C** type components (capacitors, chemical species), **R** type components (resistors, reactions), **I** type components (inductors, point masses), and energy sources (batteries, chemostats). For more examples and details of bond graph components, refer to Gawthrop and Bevan (2007) and Borutzky (2011).

Components are joined together with a bond, which represents the energy transfer between components. Each bond has an effort and flow variable associated with it, and a direction which determines sign convention. Bonds may also connect to junctions, which describe particular conservation laws. These are either *Equal Effort* (effort is equal in connected components) or *Equal Flow* (flow is identical). In bond graph notation, these are equivalent to the **0** and **1** junctions respectively.

Since bonds are two-way energy connectors, the energy of the entire system is always conserved. (Assuming a closed system. In an open system, energy can be dissipated or added, but it can always be tracked.) A similar rule applies to mass conservation or charge conservation. Bond graphs therefore by design enforce physical and thermodynamic constraints.

### B. Model descriptions

Details of example models used in Sections 3.1 and 3.2.

#### B.1. Ion Transport

Parameters and Initial Conditions

**Table 2.** Internal and external concentrations and charges for the ions Na^+^, K^+^, and Cl ^−^. Taken from Cudmore et al. (2021).

| Ion | charge $z$ | $c_{in}$ [M] | $c_{ex}$ [M] |
| --- | --- | --- | --- |
| Na<sup>+</sup> | +1 | 0.019 | 0.155 |
| K<sup>+</sup> | +1 | 0.136 | 0.005 |
| Cl<sup>-</sup> | -1 | 0.078 | 0.112 |

#### B.2. SERCA

Chemical Equations

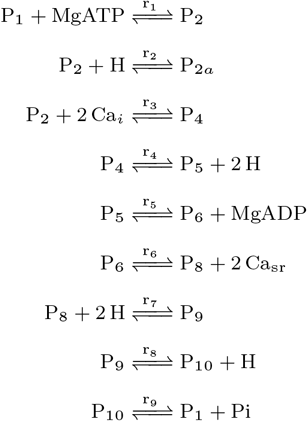

Parameters and Initial Conditions

**Table 3.** Parameter values and initial conditions for the states used in the SERCA bond graph model. Taken from Pan et al. (2020). The input function for Casr was chosen to resemble a physiological calcium spike, which is characterised by an initial spike in concentration of Ca^2+^ followed by an exponential decay.

| Reaction | Rate [s<sup>-1</sup>] |
| --- | --- |
| r<sub>1</sub> | 0.00053004 |
| r<sub>2</sub> | 8326784.0537 |
| r<sub>3</sub> | 1567.7476 |
| r<sub>4</sub> | 1567.7476 |
| r<sub>5</sub> | 3063.4006 |
| r<sub>6</sub> | 130852.3839 |
| r<sub>7</sub> | 11612934.8748 |
| r<sub>8</sub> | 11612934.8748 |
| r<sub>9</sub> | 0.049926 |

| Species | $x(t=0)$ [mol] |
| --- | --- |
| P1 | 0.000483061870385487 |
| P2 | 0.0574915174273067 |
| P2a | 0.527445119834607 |
| P4 | 1.51818391164022e-09 |
| P5 | 0.000521923287622898 |
| P6 | 7.80721128535043e-05 |
| P8 | 0.156693953834181 |
| P9 | 0.149232225342376 |
| P10 | 0.108044124948978 |

| Species | Affinity [mol<sup>-1</sup>] |
| --- | --- |
| P1 | 5263.6085 |
| P2 | 3803.6518 |
| P2a | 3110.4445 |
| P4 | 16520516.1239 |
| P5 | 0.82914 |
| P6 | 993148.433 |
| P8 | 37.7379 |
| P9 | 2230.2717 |
| P10 | 410.6048 |
| Cai | 1.9058 |
| Casr | 31.764 |
| MgATP | 244.3021 |
| MgADP | 5.8126e-7 |
| Pi | 0.014921 |
| H | 1862.5406 |

**Table 3.** Parameter values and initial conditions for the states used in the SERCA bond graph model.
| Chemostat | Concentration [mol] |
| --- | --- |
| Cai | 0.0057 |
| H | 0.004028 |
| MgADP | 1.3794 |
| MgATP | 3.8 |
| Pi | 570 |
| Casr | $\frac{2.5 \sin(0.2t)}{(t^2+0.5)} + 0.2 \exp\left(\frac{-t}{100}\right)$ |

### C. Julia Code

Full Julia code used to create the bond graph models and figures used in the main article and supplementary material. Output equations and plots (Figure 1) are labelled with #OUTPUT.

#### C.1. Ion pore transport bond graph model

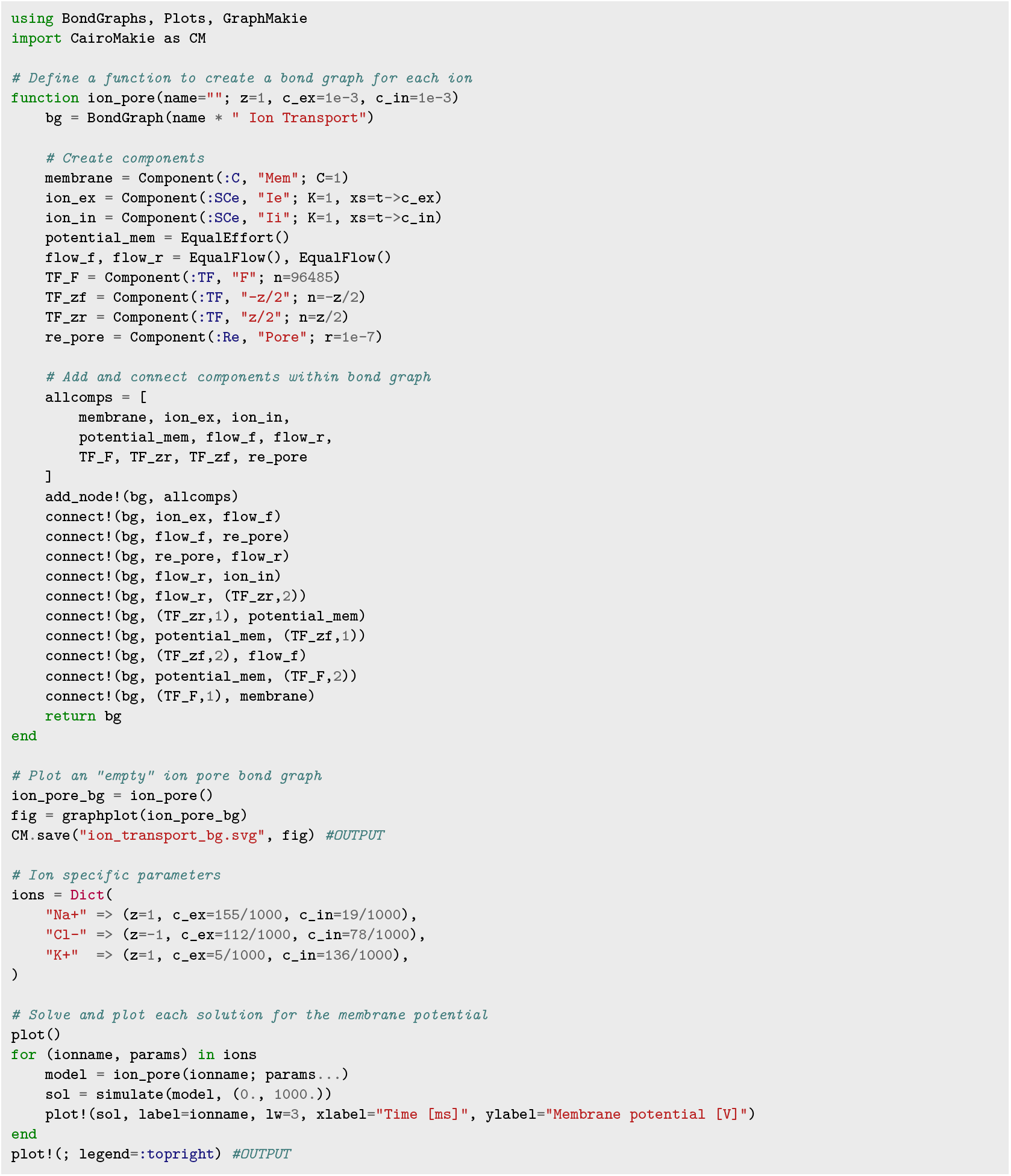

#### C.2. SERCA bond graph model

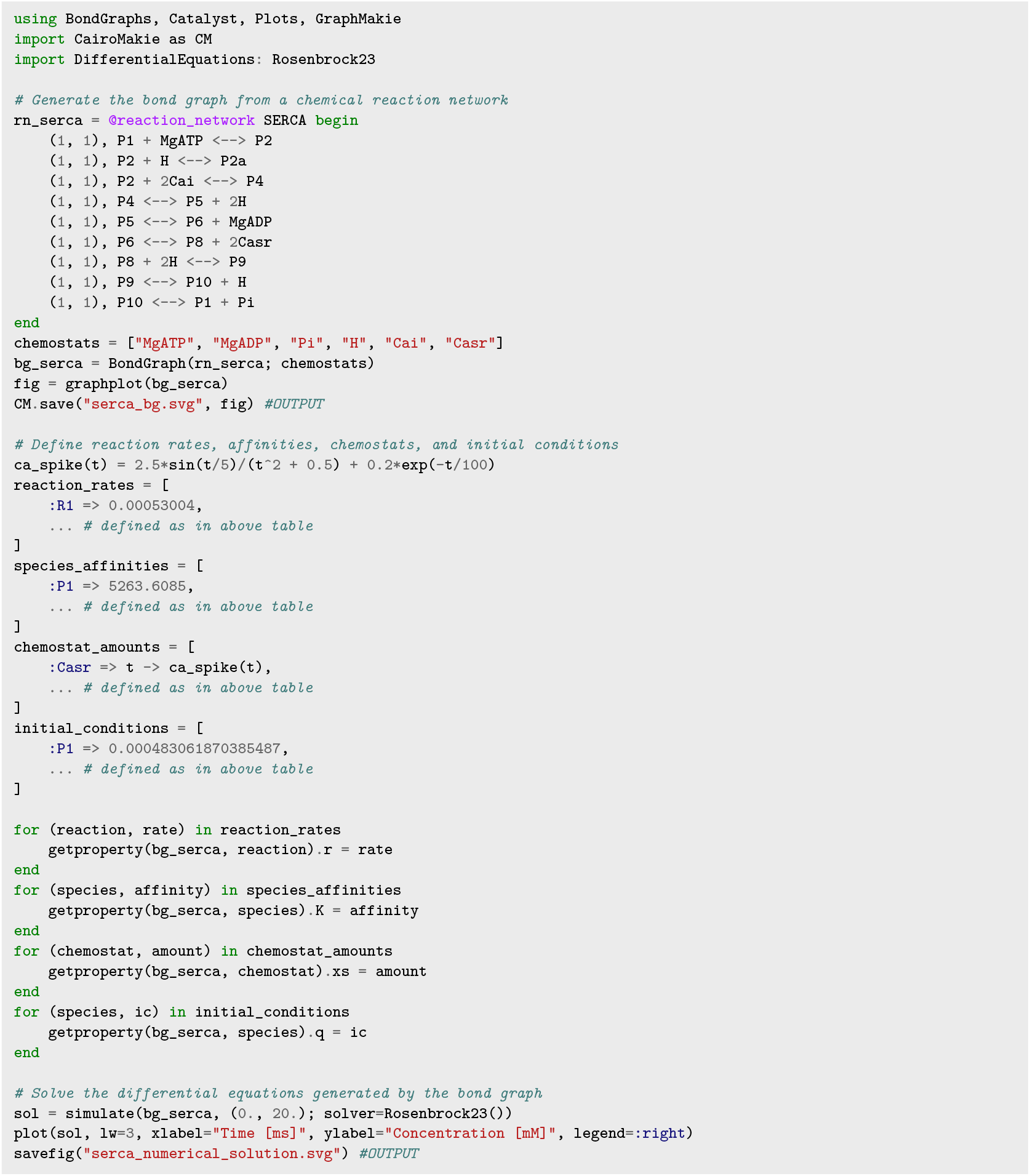

## Footnotes

1 https://jedforrest.github.io/BondGraphs.jl/stable/

